# Cross-modal quality transfer: enhancing MEG spatial resolution using BOLD-fMRI and Explainable machine learning

**DOI:** 10.64898/2025.12.16.688826

**Authors:** Jiri Benacek, Krish D. Singh, Derek K. Jones, Phoebe Asquith, David Marshall, Simon K. Rushton, Marco Palombo

## Abstract

**Purpose:1:** Blood-oxygen-level-dependent functional MRI (BOLD-fMRI) and magnetoencephalography (MEG) offer complementary insights into brain function, with BOLD-fMRI providing high spatial resolution and MEG offering high temporal sensitivity. This study aimed to enhance MEG’s spatial resolution through learning inter-modal relationships via data-driven fusion with BOLD-fMRI using explainable machine learning (xML).

**Methods:** MEG and BOLD-fMRI data were collected for sixteen participants watching the same naturalistic visual stimulus. MEG signals were filtered into standard frequency bands and processed to match the haemodynamic response. Data were then decomposed using Tensorial Independent Component Analysis (TICA), yielding 250 MEG (25 per frequency band) and 30 BOLD-fMRI components. A set of Extreme Gradient Boosted tree (XGBoost) models were trained to predict MEG component activity from downsampled BOLD-fMRI components. The models with best-performing configuration of hyperparameters at this resolution were then used to generate voxelwise upsampled MEG maps using the natively higher-resolution BOLD-fMRI data. Performance was compared to trilinear interpolation using R², mean squared error (MSE), and structural similarity index (SSI). Model interpretability was enhanced by generating Shapley values (SHAP), describing relationships between input data and model output.

**Results:** Models trained on 6mm³ data achieved high predictive performance on previously unseen data (R² = 0.80, MSE = .07). Using higher-resolution 2mm³ BOLD-fMRI data, MEG activity was upsampled to 2mm³, resulting in more detailed image, while maintaining reasonable congruence to naive interpolation (R² = .81, MSE = .06).

**Conclusions:** This work demonstrates that explainable machine learning enables spatial super-resolution of MEG maps via BOLD-fMRI-informed quality transfer, enabling enhancement of spatial resolution, and offering improved localization, as well as means of generating data-driven insights into neurovascular coupling.

## 1. Introduction

Functional magnetic resonance imaging (BOLD-fMRI) and magnetoencephalography (MEG) each have unique strengths and limitations. MEG provides measurements that directly reflect magnetic fields generated by the electrical activity of neuronal populations, but it has low spatial resolution and reduced sensitivity to subcortical signals [21]. BOLD-fMRI provides finer spatial resolution with whole-brain coverage, but only indirectly through blood-oxygen-level-dependent signal (BOLD).

Combining BOLD-fMRI and MEG data is an attractive avenue to leverage their complementarity, but the different origin of their signals presents challenges. BOLD responses arise from a complex interplay between energetic demands and processes such as glucose metabolism [51] and oxygen extraction fraction [7] and reflect changes in blood oxygenation levels coupled with neural activity, rather than the activity itself. Interpreting BOLD-fMRI signals is challenging due to limited understanding of the exact neurophysiological underpinnings and their relationship to neuronal firing rates and various components of oscillatory local-field potentials, encompassing both excitatory and inhibitory processes. [28, 48, 12].

Published literature on neurovascular coupling has shown that BOLD signals are often closely related to neuro-electric activation, with the strongest temporal correlations observed between BOLD and oscillatory activity in the gamma frequency range above 30Hz [29]. However, this relationship becomes more complex at lower frequencies, where the coupling is negative [36, 9], and even nonlinear [11, 13, 20]. Although some research has combined imaging modalities such as BOLD-fMRI and EEG, they often prioritize time-domain analyses while overlooking the possibility that relationships may vary across different frequency bands. [53].

There have been previous efforts to combine MEG and BOLD-fMRI, aiming to leverage the complementary strengths of the modalities. Early work focused on integrating MEG’s temporal precision with BOLD-fMRI’s spatial resolution through constrained source localisation, in which BOLD-fMRI-derived activation maps were used as priors to guide MEG inverse modelling [40, 10]. Newer approaches have extended beyond source reconstruction to data-driven fusion methods, including joint independent component analysis [33] and canonical correlation analysis [57], which identify shared patterns of spatial or temporal variability across modalities. More recently, generative models including Bayesian and dynamic causal modelling approaches have also incorporated priors, using BOLD-fMRI to inform source localization and neurovascular coupling models in MEG [22], and novel techniques such as high spatial-frequency component extraction from BOLD-fMRI have been leveraged to enhance MEG/EEG source localisation within the normally inaccessible deep regions [34]. These methods have provided insights into the overlap and divergence of MEG and BOLD-fMRI functional networks, but typically assume linear coupling and do not readily capture spatially heterogeneous or complex relationships. A smaller body of work has explored the inverse direction — using MEG to inform BOLD-fMRI analysis — for example, to improve detection of transient events in naturalistic paradigms [6]. However, few studies have investigated whether the more spatially precise BOLD-fMRI can be used to directly reconstruct high-resolution representations in the less spatially sensitive MEG, particularly in a fully data-driven framework without the need for imposing a priori coupling assumptions.

In this work, we propose a novel, data-driven approach to improve the spatial resolution of MEG-based brain activity mapping by leveraging complementary information from BOLD-fMRI, and learn the relationships between MEG and BOLD-fMRI signals’ independent components to generate high-resolution MEG maps from BOLD-fMRI inputs using explainable machine learning (xML, i.e. ML models with transparent decision making). We believe that if the model succeeds in learning strongly predictive cross-modal relationships, the explainability of our results can also provide a tentative step towards a more comprehensive understanding of the interplay of the two functional imaging approaches without imposing a priori hypothesis, thanks to its purely data-driven nature. To investgate this, we used a unique dataset of matched multimodal data with participants undergoing BOLD-fMRI and MEG scans while watching the same movie clip.

## 2. Methods

Our objective is to develop a data-driven machine learning model that can learn the relationship between the components of both MEG and BOLD-fMRI signals and use the learned relationships to subsequently predict high spatial resolution MEG components from BOLD-fMRI input. To enable this, we will use the naturalistic viewing paradigm to synchronize the brain activity between the two modalities, as well as across scans, in a cohort of healthy volunteers. In this section we describe the cohort, the rationale for using the naturalistic viewing paradigm and what machine learning model and explainability techniques we used.

### 2.1. Participants

16 healthy volunteers (n=9/7 M/F; average age 27) with normal vision and no history of neurological disorders underwent both MEG and BOLD-fMRI scans. All subjects gave informed consent to take part in the study and ethical approval was granted by Cardiff University School of Psychology Ethics Committee. To reduce or eliminate systematic bias, half of the subjects had watched the same movie clip within the MEG scanner, before watching it within the MR scanner, and half watched the clip in MR first before moving to MEG, with the two acquisitions happening maximum 5 months apart.

### 2.2. Paradigm

Data was collected using naturalistic viewing paradigm where participants were instructed to watch a movie as they normally would (i.e. without any imposed restrictions), following work of Bartels and Zeki (2004) [4]. This paradigm offers a more ecologically valid way of measuring functional activity by presenting rich, dynamic and engaging stimuli, while preserving strengths of task-based and resting-state approaches [1, 50]. In BOLD-fMRI they show highly replicable activation patterns on both voxel- [19] and feature-level [4], also showing a good ability to predict task-based brain activation maps [17], with high test-retest reliability [50]. Similarly, in MEG, time-locked analyses during naturalistic viewing have demonstrated robust and spatially consistent activation and connectivity patterns highly concordant with classical fMRI results [38], with good evidence of inter-subject consistency in the temporal domain [26], and even similarities between envelopes of MEG canonical variates and fMRI voxel time-courses [27].

In both, MEG and BOLD-fMRI, participants watched a 19 min 40 sec movie clip taken from the opening sequence of Skyfall (2012) [54], edited to ensure it featured a broad range of stimuli such as faces, buildings, machines, text, objects, social interactions, as well as camera movements and a strong narrative to hold participants’ attention. Subjects were told to watch the clip as they would normally do, with no restrictions on fixation.

### 2.3. Imaging

#### 2.3.1. Functional Magnetic Resonance Imaging (BOLD-fMRI)

##### Data acquisition

BOLD responses were acquired with a 3T General Electric (GE) Signa 3T HDxt Scanner (General Electric, Milwaukee, Wisconsin), equipped with echo-speed gradient coil and amplifier hardware using a GE 8 channel receiver RF coil. Activation images were acquired using echo-planar imaging (EPI) with a TR of 2 seconds, TE of 30ms, flip angle of 76^◦^(Ernst angle calculated), and with spatial resolution (3*mm*^3^), later registered to common space at 2*mm*^3^ and 6*mm*^3^ to allow for comparative analyses. At every TR a volume of 37 contiguous slices was acquired to cover the whole brain. Slices were aligned to the anterior-posterior line of the participants’ corpus-callosum. High-resolution anatomical scans were collected for each participant (resolution 1*mm*^3^), using a fast-spoiled gradient echo scan sequence (FSPGR)

##### Data processing

Pre-processing of the BOLD-fMRI data for was performed in FSL version 6.0.5. [24] via BOLD-fMRI Expert Analysis Tool (FEAT) using default settings: data were smoothed at 5mm; motion corrected using FSL’s Linear Image Registration Tool (mcFLIRT) [23] ; and highpass temporal filtering at 100Hz was used to remove low frequency temporal drifts.

Each subject’s EPI data were extracted using BET, and then registered to their own FSPGR using FMRIB’s Linear Image Registration Tool (FLIRT) [25], with either, a 6DoF tranform (n=5), or using Brain-Boundary registration (n=11), chosen based on visual inspection. After the functional data were registered to participant’s own structural image, the data was registered to standard space (MNI152[15]) using FNIRT’s (FMRIB’s nonlinear Image Registration Tool). Later downsampling onto 6mm to spatially match the MEG was performed using the FSL ‘applywarp’ command. The processing pipeline is shown in Fig.1.

**Figure 1:**
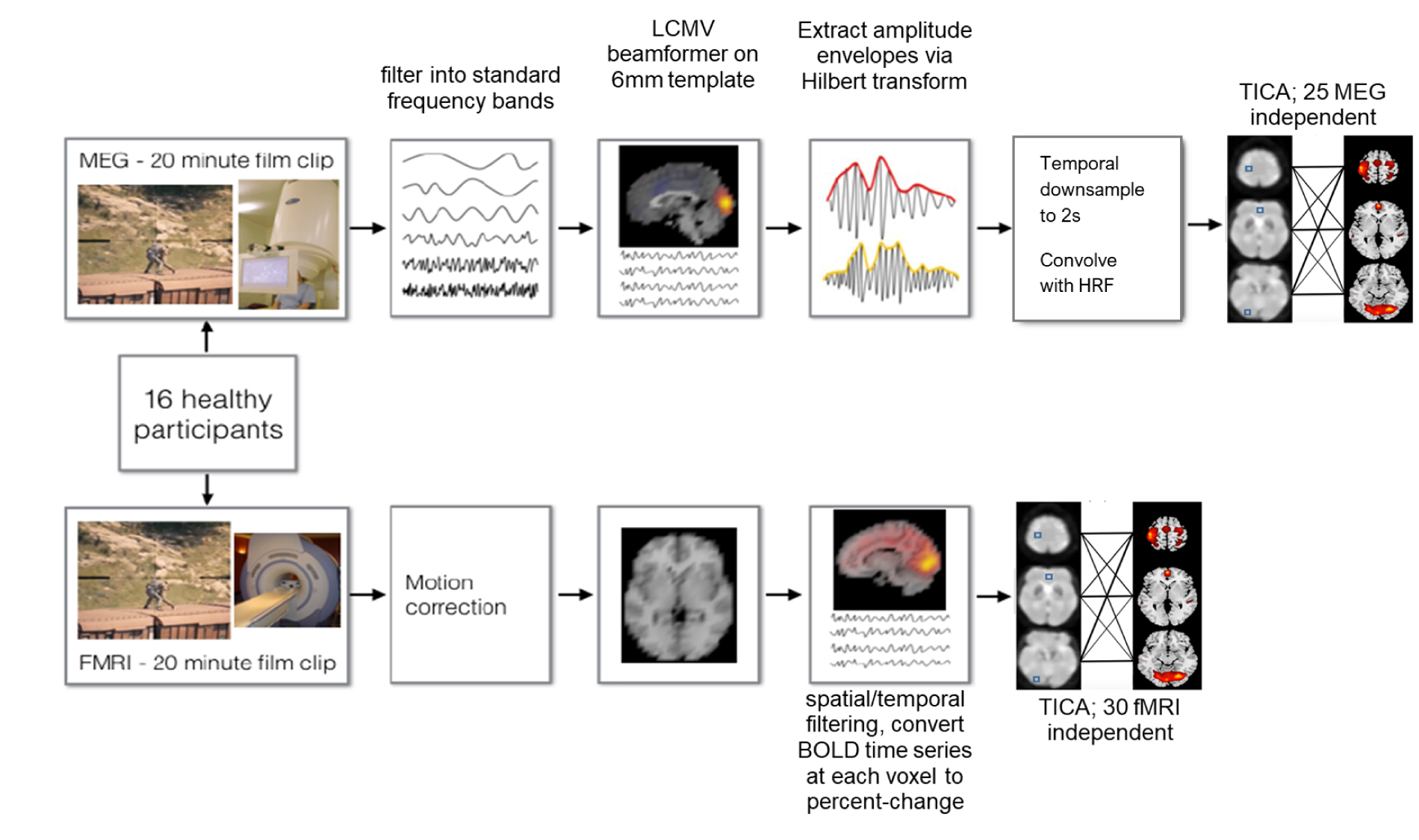
Pre-processing pipelines for BOLD-fMRI and MEG. Figure adapted from [43, Lu]

#### 2.3.2. Magnetoencephalography (MEG)

##### Data acquisition

Recordings were collected using a CTF Omega 275-channel MEG system sampled at 1200Hz and analysed as synthetic third-order gradiometers. Continuous head localisation of electromagnetic coils placed at fixed distances from anatomical landmarkss relative to anatomical landmarks (10mm anterior to left and right tragus, 10mm superior to nasion) was then used for MEG/MRI co-registration. Fiduciary locations were manually marked onto the anatomical MRI after the scanning session.

##### Data processing

Pre-processing was done using a combination of FieldTrip [39] and SPM12 [2]. Data were down-sampled to 600Hz. Predefined functionally-specific frequency bands were used to filter the raw data: delta (*δ*; 1-4Hz), theta (*θ*; 4-8Hz), alpha (*α*; 8-13Hz), beta (*β*; 13-30Hz), and low-gamma (low-*γ*; 40-60Hz), and a collection of high-gamma bands (high-*γ*; 60-80, 80-100, 100-120, 120-140, 140-149). Data at each frequency band were source localized using a linearly-constrained minimum-variance (LCMV) beamformer Synthetic-aperture magnetometry (SAM) onto an MNI152-template space at a resolution of 6*mm*^3^, and downsampled to 2s to match the BOLD-fMRI readings. This provided us with virtual time-series at each voxel in the brain. Using the Hilbert Transform and analytic function, time-series were converted to an amplitude envelope which was de-spiked and high-pass filtered at *>*0.01Hz, removing drift. Due to lack of regional estimates for the HRF, the processed MEG data were then convolved with the canonical HRF from SPM to match the expected BOLD response. This was performed to allow a direct comparison between the MEG and MR data collected during the movie watching. The processing pipeline is shown in Fig.1.

#### 2.3.3. Feature extraction

For feature extraction, a group-level Tensorial Independent Component Analysis (TICA) [5] from FSL version 6.0.5. [24] was applied separately to the BOLD-fMRI and MEG data, yielding 30 BOLD-fMRI independent components (ICs) and 25 MEG ICs per frequency band, all in a shared MNI152 space. The range of extracted components was chosen to reflect previous literature, which often uses ∼20 components to clearly distinguish known functional networks at high level [42]. We opted for a slightly higher number of BOLD-fMRI components than MEG components to increase the likelihood of capturing sub-networks within broader networks, thereby providing a richer and more granular feature set for use as predictive input.

The resulting components have then been correlated to existing resting state network (RSN) atlas [49]. These correlational analyses showed that components extracted from our naturalistic viewing BOLD-fMRI data match common RSNs (See Supporting information Figure A1), with most visual networks being well-represented by one or two components BOLD-fMRI, but being highly non-specific in MEG across the extracted frequency bands (See Supporting information Figure A2, a-e), suggesting highly complex and nonlinear relationships between modalities.

#### 2.3.4. Feasibility study: Recovering maps using classical curve fitting

Prior to machine learning modelling, we conducted a feasibility study to determine whether it is possible to learn MEG ICs from BOLD-fMRI ICs using classical optimisation techniques and whether this representation remains stable enough to facilitate upsampling through quality transfer (e.g. enhancing spatially low-resolution MEG data by using high-resolution BOLD-fMRI data).

To test this, we trained an L2 regularised linear model on 6*mm*^3^ data, using MEG ICs extracted across frequency bands to predict each BOLD-fMRI IC. Following training, we applied nonlinear curve fitting to solve the inverse problem, (i.e. from BOLD-fMRI ICs to MEG ICs) using Powell’s algorithm constrained within the minimum and maximum activation bounds of the original (6*mm*^3^) MEG data, with increasingly more noisy initial guess provided by an interpolated version of said MEG ICs (6*mm*^3^), with normally distributed noise levels set at 0, 0.01, 0.05, 0.1, and 0.2.

The algorithm was then used to optimize the interpolated MEG inputs for best-fitting of an axial slice from a 2*mm*^3^ BOLD-fMRI map of early visual network (Component 7) by minimizing its mean squared error (MSE) with the model’s prediction, aiming to recover each MEG IC at a higher-than-trained resolution from the pre-trained model. The optimisation model’s schematic representation can be found at 2a, along with examples of recovered maps 2b-2d, as well as results of this experiment in Table in 2e.

When initialized using the interpolated values, the Powell’s optimization recovered did not strongly deviate from the initial guess, with MSE ≤ 0.01, MAE ≤ 0.01, and *R*^2^ = .99. However, when initialized with noise-injected interpolated images, the MEG ICs recovered after the Powell’s optimization were increasingly further from the interpolation baseline (MSE = 0.93, MAE = 0.13, and *R*^2^ = 0.56 at 0.2 noise level); suggesting a complex loss landscape with multiple local minima. Thus, we explored whether modern machine learning can stabilize this mapping.

#### 2.3.5. MEG up-sampling: modelling and analysis

To predict MEG ICs from BOLD-fMRI ICs, we employed gradient boosting ensemble models, as they don’t require explicit regularization, can learn data-driven representations and priors, and capture nonlinear relationships between features and targets that classical linear optimization cannot [16]. In our design, we carefully controlled for overfitting while maintaining interpretability by using cross-validation.

A series of regression models (N=250 in total: one for each of the 25 MEG components and 5 frequency bands) were trained to predict each MEG component/bands from the 30 BOLD-fMRI components downsampled to match MEG’s resolution (6*mm*^3^), and the best-performing models were subsequently used for upsampling MEG’s spatial resolution at voxel-level by inputting the 30 non-downsampled (2*mm*^3^) higher-resolution BOLD-fMRI components. We chose to utilise a tree-based Extreme Gradient Boosting (XGBoost) [8] because of its good combination of predictive power, and explainability. XGBoost builds the final model iteratively by adding trees sequentially, where each new tree learns to correct the prediction errors made by the ensemble of previously constructed trees. We split the data into training and testing subsets, used 3-fold cross-validation on the training subset for modelling to systematically test all combinations of algorithm settings (hyperparameters) using grid search to find the optimal values. Tuned parameters included the number of estimators (1 to 250), learning rate (0.05 to 0.23), and tree depth (1, 2, or 5). We built the final model using the entire training dataset with the parameter settings that performed best during cross-validation, and then evaluated it on the unseen data in held-out test set. To further enhance the explainability, local explanations (e.g. explanations for each prediction) using Shapley additive explanations (SHAP) [31] were applied to quantify how strongly each input feature (BOLD-fMRI component) influenced the prediction and whether that influence was positive or negative with respect to the output (predicted/upsampled MEG component).

Results were evaluated in terms of *R*^2^, MSE, and SSI [52]. First, we assess the quality of the mapping by comparing the predictions against ground truth at baseline 6*mm*^3^, and, subsequently, in absence of a true ground-truth, we compare the predicted 2*mm*^3^ maps to the baseline maps upsampled through trilinear interpolation using FSL’s ’applywarp’, in order to investigate robustness of the model, and to highlight anomalies in the prediction such as hallucinations. A schematic of the implemented approach is shown in Fig. 3.

**Figure 2:**
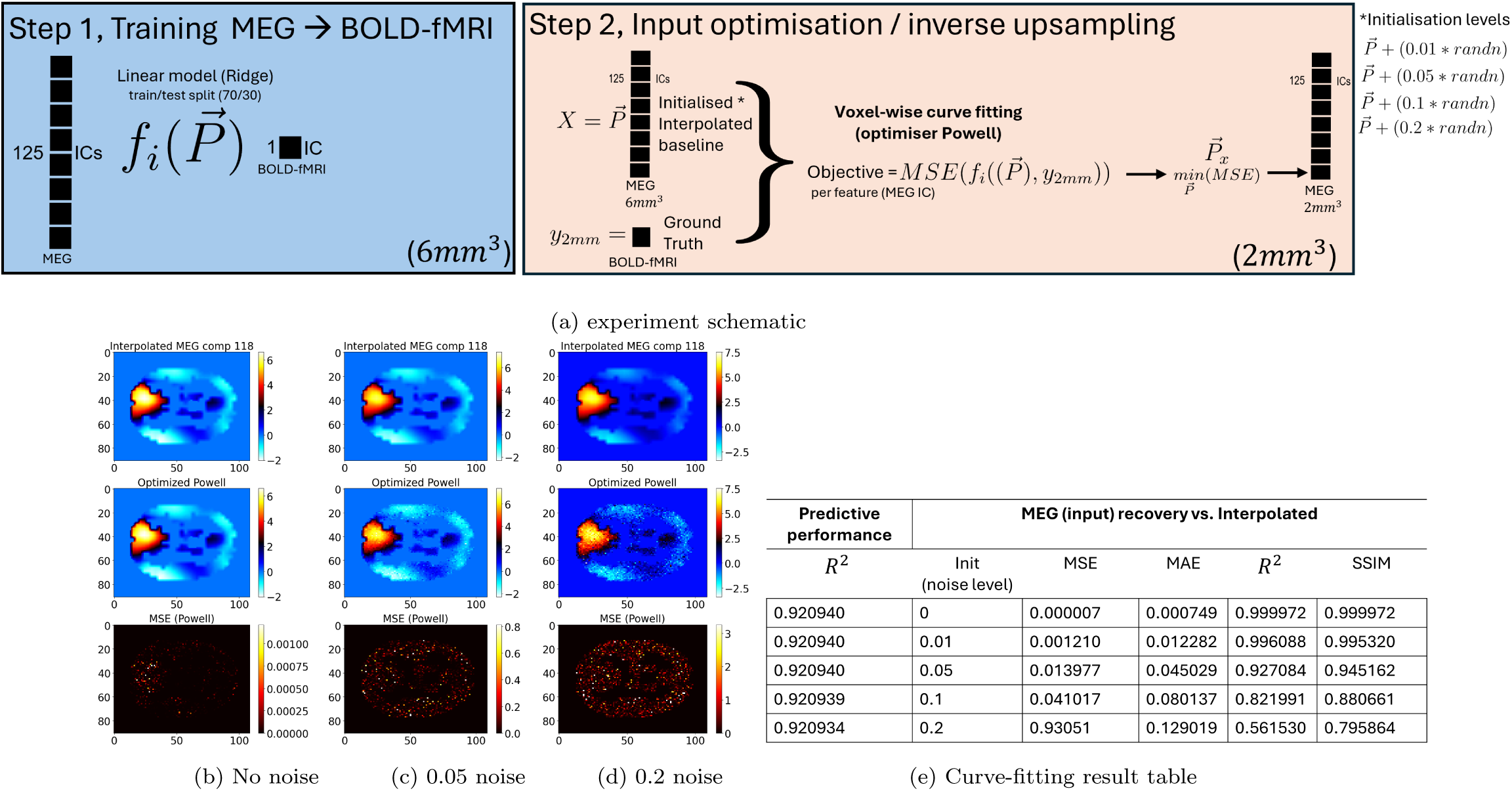
a) Powell voxel-wise input optimisation experiment’s schematic representation.; **b)-d)** various levels of noise-injected interpolation initialisation recovery of MEG Component *low* − *γ*18. From top: Interpolation, Powel-optimisation, Error map (MSE). MSE levels have been thresholded to 25% of the maximum value for better visibility; **e)** Prediction & Recovery performance

**Figure 3:**
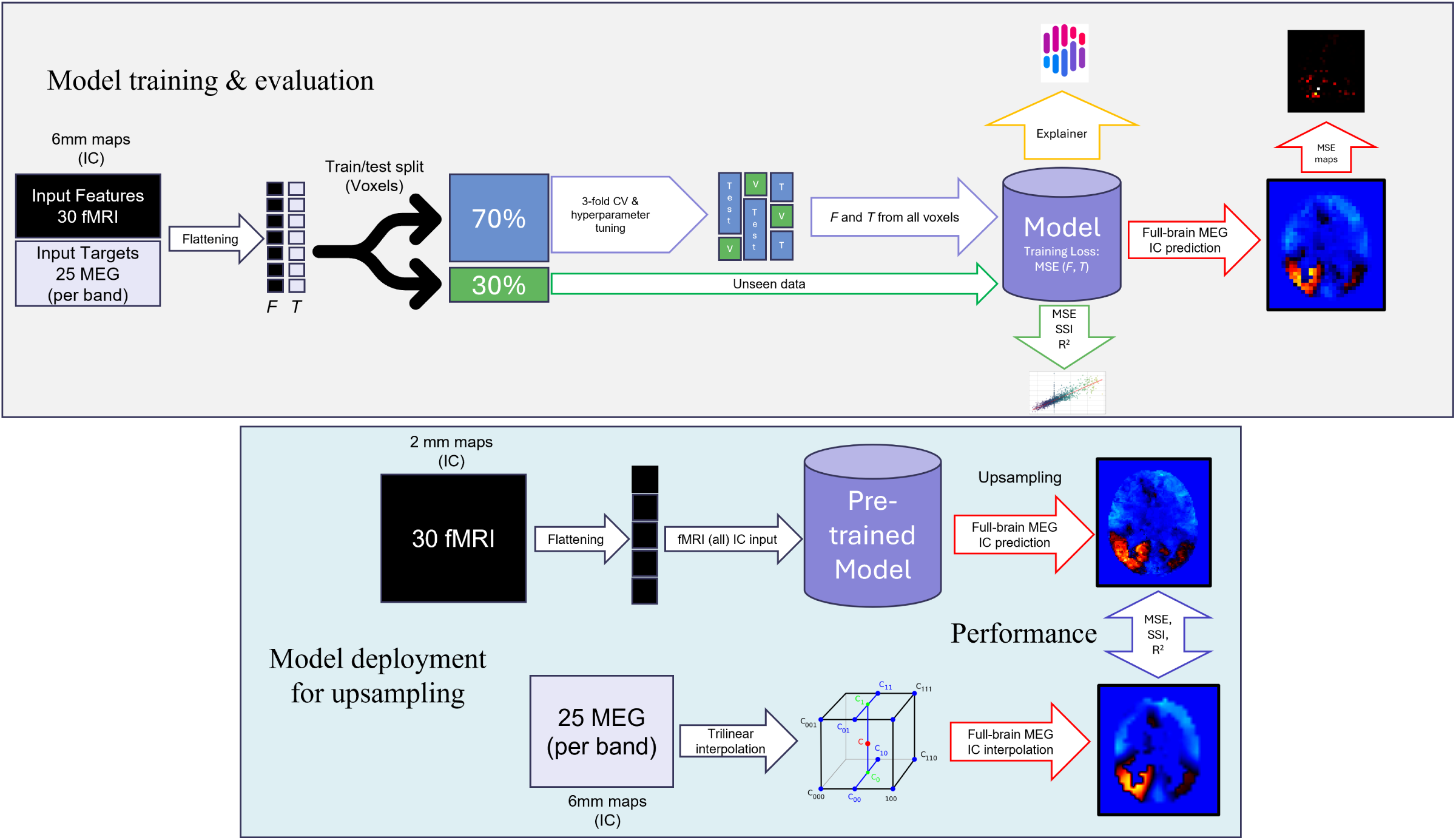
Diagram of the model-training pipeline & model deployment for upsampling. The flattened data were split into train (70%) & test(30%) subsets, best performing set of hyperparameters was chosen using a 3-fold cross-validation on training data, and peformance was subsequently evaluated on unseen test subset. Model training has been done using downsampled (6*mm*^3^) fMRI maps as input features for voxel-wise prediction of MEG component at a matching resolution. The deployment to higher resolution’s performance compared MEG maps upsampled on voxel-level using the previously trained model, with input fMRI features at high resolution (2*mm*^3^) against a naive baseline of MEG data interpolated to 2*mm*^3^. F = Feature, T = Target

## 3. Results

### 3.1. Upsampling Early visual networks

Because of strong evidence of good congruence between MEG gamma-band and the BOLD response in primary visual cortex [37], results were generated for component 18 map from low-*γ* (40-60Hz) band, positively (r = .43) correlated with early visual areas as described in [49].

#### 3.1.1. Upsampling using ML

We trained the set of models on 6*mm*^3^ data, using BOLD-fMRI signals to predict each of the MEG components across frequency bands. After training, we inputted 2*mm*^3^ BOLD-fMRI data and predicted MEG components in the upsampled 2*mm*^3^ spatial resolution. The final model achieved *R*^2^ = .80 for unseen test subset at 6*mm*^3^ (*R*^2^ = .92 for full data), and *R*^2^=0.81 for full image at 2*mm*^3^ when compared with a reference of naively interpolated map, used as reference in absence of a true ground-truth (Fig 4).

**Figure 4:**
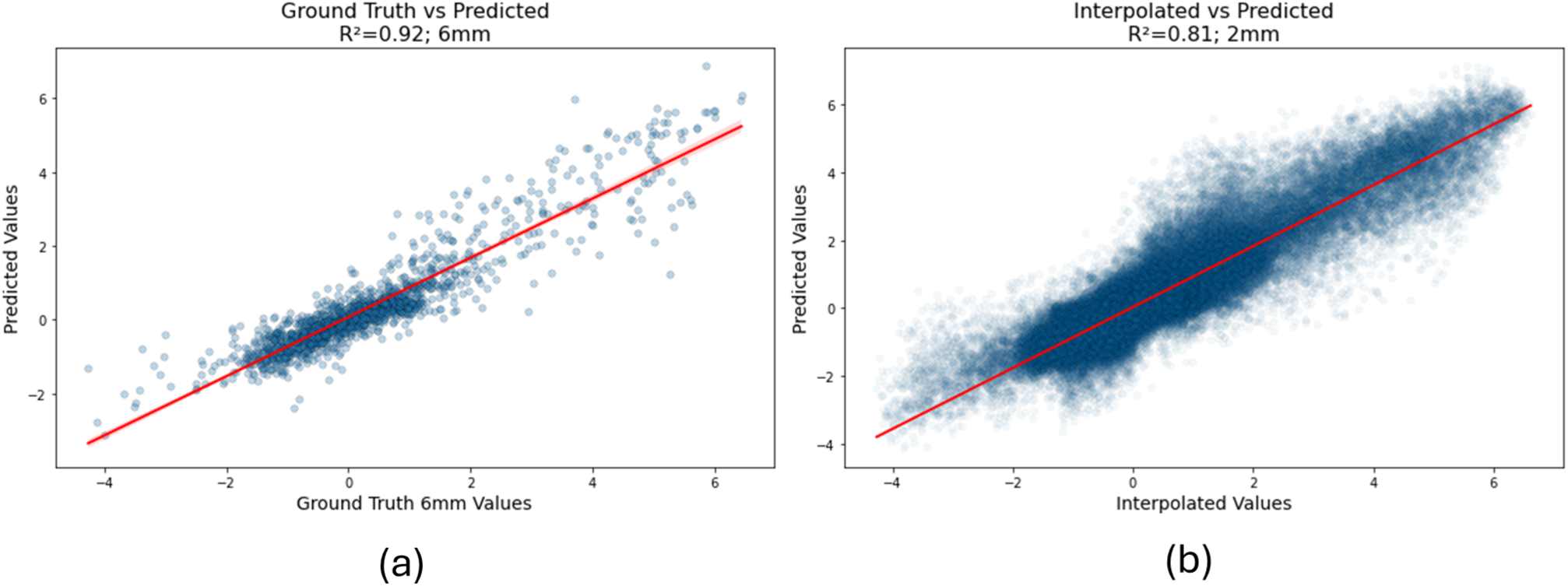
**(a)**: Scatterplot for model performance for Test subset at 6mm with ground truth values for low-*γ* 18 (x-axis) and model predictions (y-axis), *R*^2^ = .92. **(b)** Scatterplot for model performance for full data at 2mm with trilinearly interpolated values for low-*γ* 18 (x-axis) and model predictions (y-axis), *R*^2^ = .81.

Results are presented in Fig.5a-c. Fig.5a displays the spatial maps and deviations be-tween the maps at 6*mm*^3^ resolution (MSE = .030; SSI = .93). Maps remained robust when compared to the interpolation to 2*mm*^3^ (MSE = .063; SSI = .86). Fig.5b highlights that the model is strongly influenced by the highly correlated BOLD-fMRI component 7 (Fig.5c), while being modulated by less predictive components with both positive and negative relationships with the predicted upsampled MEG component.

The resulting maps predicted by the model were visually inspected to confirm the absence of any apparent artifacts or hallucinations. The model-upsampled maps demonstrated greater detail compared to those generated through interpolation alone. Notably, the maps upsampled by our method show emergent patterns not present in MEG that resemble brain’s structural details, suggesting that these patterns may be encoded within the BOLD signal and transferred via modelling.

### 3.2. Upsampling Delta 23

To examine and contrast how the method performs in areas where we saw less direct relationship between the modalities, we then predicted maps for component *δ*23 (Fig. 6), partially correlated with BOLD-fMRI-derived component corresponding to default mode network (r=.12), cerebellum (r=.12), and visual areas (r=.18) [49]. While at 6mm, we saw similar performance to low-*γ*18, with MSE = .05, SSI = .93 and *R*^2^ = .89, at 2*mm*^3^, the model was less similar to its counterpart derived via interpolation (MSE = .12, SSI = .84, *R*^2^ = .69). SHAP analysis (Fig. 6b) revealed negatively directed relationships with the top predictors, with different component maps contributing more evenly then in the case of low-*γ*18 model. However, looking at the top predictive features (Fig.6c), we can see that none directly resemble the predicted upsampled MEG component. This, combined with the lack of a dominant spatial correlate in the BOLD-fMRI input likely accounts for the lower predictive performance.

**Figure 5:**
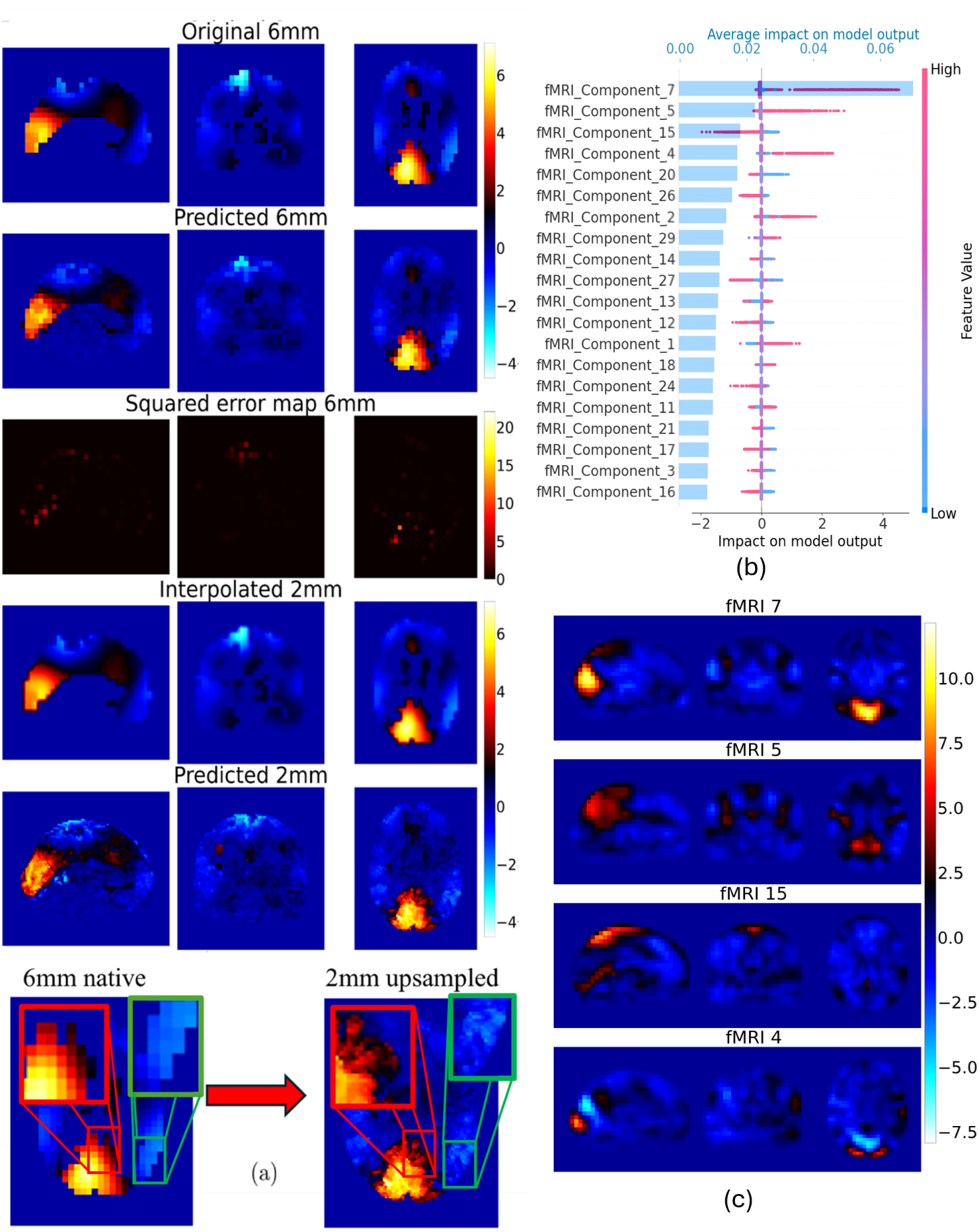
**(a)**: Component 18 from low-*γ* band (40-60Hz). From top: Ground truth 6*mm*^3^ image, 6*mm*^3^ image predicted using BOLD-fMRI input, Squared error maps between original and predicted images, interpolated 2*mm*^3^, BOLD-fMRI-guided upsampling to 2*mm*^3^, zoomed-in original 6*mm*^3^ and predicted upsampled 2*mm*^3^ (red functional activation, green emergent structural features) **(b)**: Shapley values, quantifying the predictive power of the components, where blue bars show relative component-contribution, and colour signifies directionality of the relationship. Each coloured dot represents a voxel within the BOLD-fMRI data, where the colour gradient shows the value of the voxel (red if negative, blue if positive), and the corresponding value on the *x* axis shows directionality and the impact on model output, as determined using SHAP analysis. **(c)** top 4 predictors for low-*γ* 18

**Figure 6:**
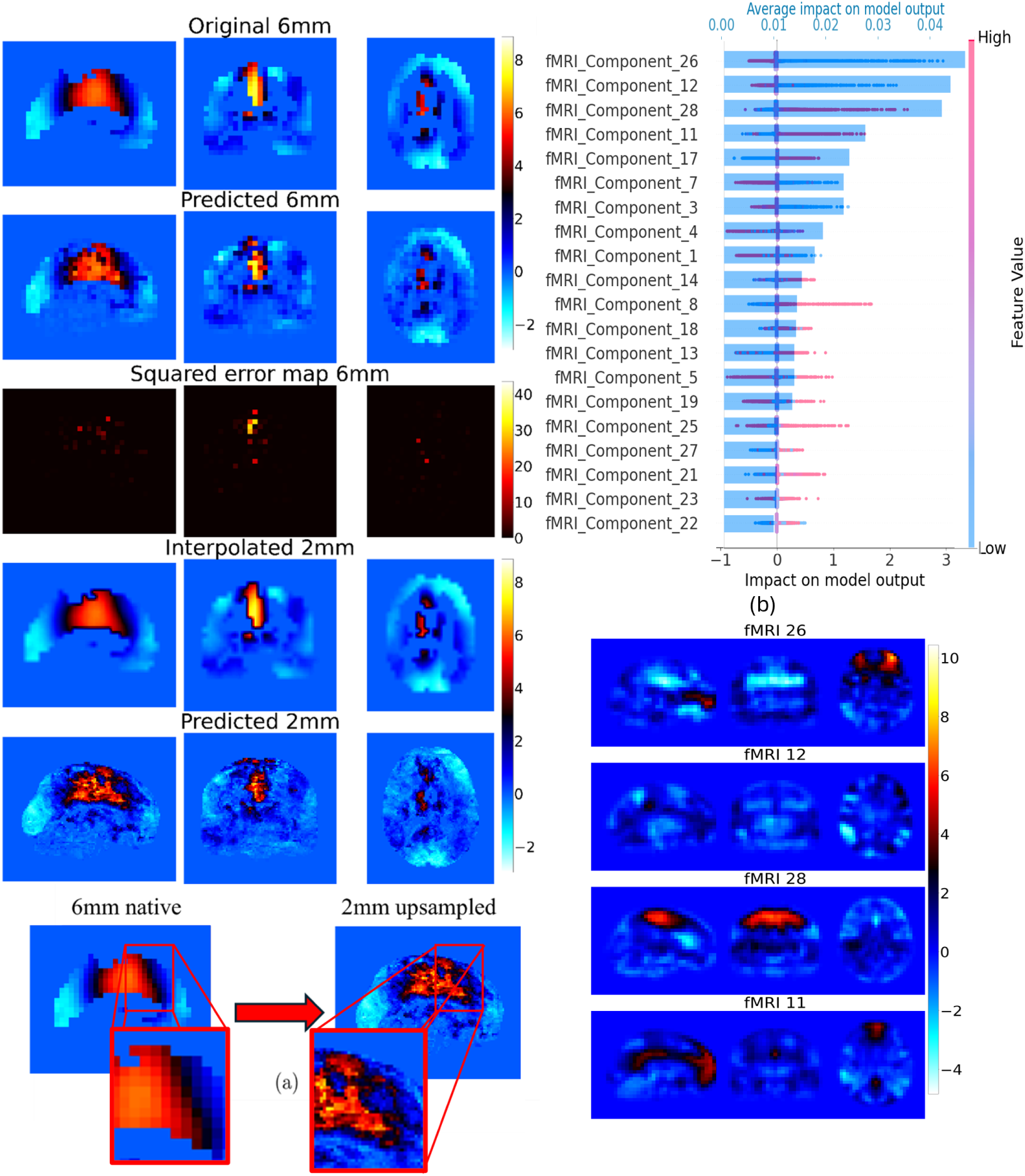
**(a)**: Component 23 from *δ* band (1-4Hz). From top: Ground truth 6*mm*^3^ image, 6*mm*^3^ image predicted using BOLD-fMRI input, Squared error maps between original and predicted images, interpolated 2*mm*^3^, BOLD-fMRI-guided upsampling to 2*mm*^3^, zoomed-in original 6*mm*^3^ and predicted upsampled 2*mm*^3^ **(b)**: Shapley values, quantifying the predictive power of the components, where blue bars show relative component-contribution, and colour signifies directionality of the relationship. Each coloured dot represents a voxel within the BOLD-fMRI data, where the colour gradient shows the value of the voxel (red if negative, blue if positive), and the corresponding value on the *x* axis shows directionality and the impact on model output, as determined using SHAP analysis. **(c)** top 4 predictors for *δ* 23

## 4. Post-hoc exploratory analyses

After the initial modelling, we performed a set of experiments to further explore SHAP explanations using UMAP (Uniform Manifold Approximation and Projection for Dimension Reduction)[32], comparison of connectomic matrices in ground truth vs upsampled models, and the emergent anatomical detail.

Additionally, we explored the model’s behaviour in terms of a) training models with denoised input, b) dealing with incomplete data, and c) stability/generalisability of the model & our feature extraction strategy (ICA) between different groups of participants.

### 4.1. Exploration of Model Explanations via SHAP-UMAP Embed-ding

To investigate latent structures in the predictive strategies employed by the models, we applied UMAP dimensionality reduction to the SHAP importance patterns across all predicted MEG components (neighbors = 5, minimal distance = 0.1, metric = euclidean) to create a 2D map where MEG components that depend on similar sets of fMRI components appear close together. The resulting embedding revealed distinct organizational features, with clusters forming for various relevant major networks/brain areas (Figure 7). Additionally, a compact cluster of components emerged, characterized by consistently low predictive performance (R²), irrespective of frequency band affiliation (Figure 7a, 7b). Upon visual inspection, we confirmed these were all noise-components.

**Figure 7:**
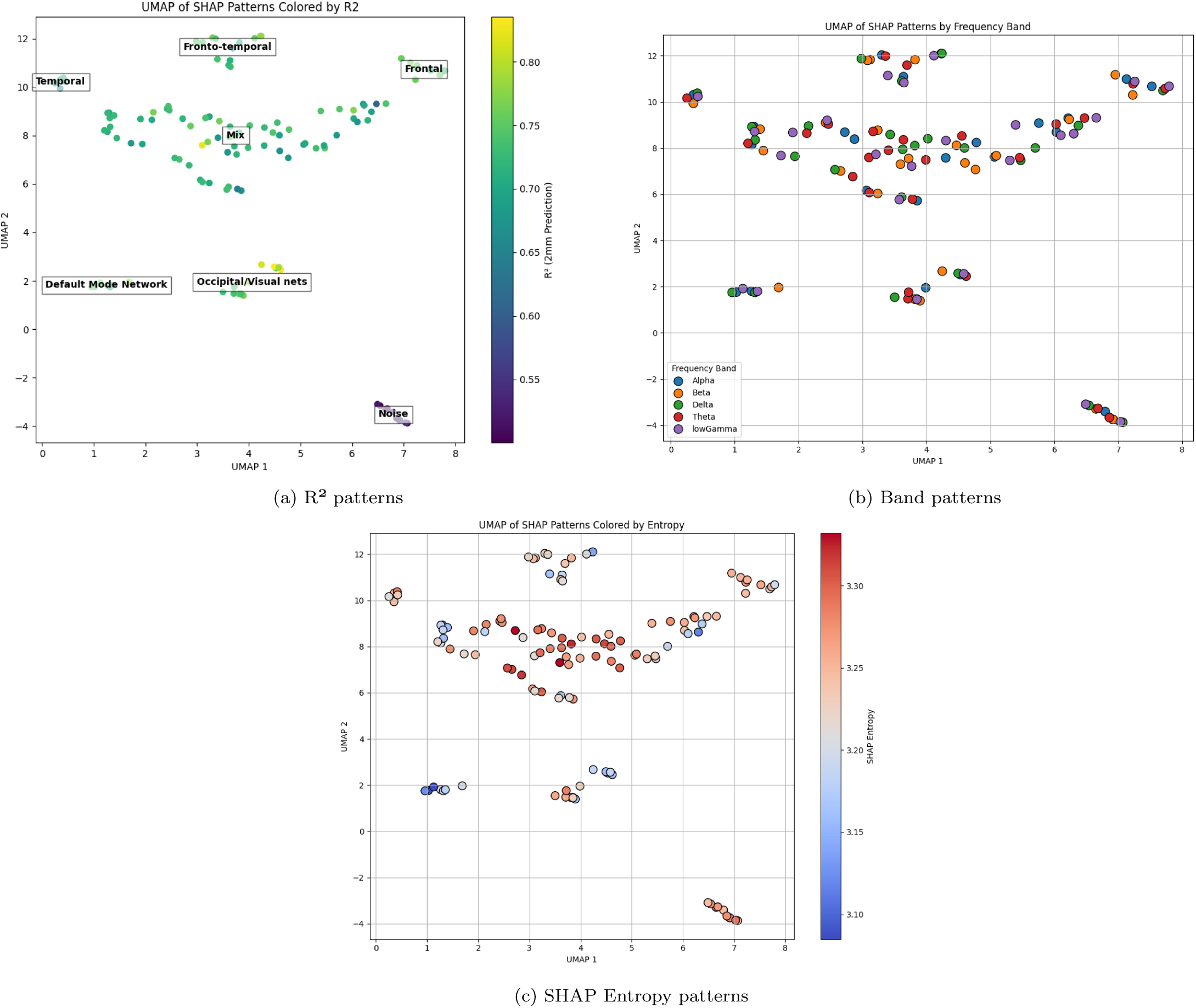
UMAP embedding of SHAP importance patterns for MEG component predictions.**(A)** Points colored by R² performance, highlighting model’s reliance on strongly-represented spatially overlapping information, as well as separating a distinct cluster of poorly-performing components (bottom-right).**(B)** Frequency band annotation reveals no segregation, suggesting shared BOLD-fMRI predictor usage across rhythms.**(C)** SHAP entropy distribution (model complexity) indicates variability in predictor focus, with no direct correlation to model success.

Next, we decided to visualise model complexity in terms of Shannon entropy [47], in which a models relying on small subsets of strong predictors would have low entropy values, and vice versa. Analysis of entropy across the SHAP embedding (Figure 7c) indicated that while some models relied on focused subsets of BOLD-fMRI predictors, others adopted more distributed strategies. Notably, focused SHAP profiles were observed both in low- and high-performing models, indicating that predictor sparsity alone does not guarantee successful reconstruction. Furthermore, the absence of frequency-specific clustering highlights a shared reliance on spatial BOLD-fMRI features across oscillatory bands, underscoring the amount of spatial cross-frequency information overlap leveraged by the models.

### 4.2. Regional Magnitude Relationships

As another means of validation, we opted to generate Regional Magnitude Relationship Matrices (Fig. 10, thresholded at |*weight*|=.25) using Schaefer et al. (2018) parcellation atlas of 100 regions of interest (ROIs) [46]. For each MEG component map, ROI-wise magnitude values were obtained by averaging voxel intensities within each parcel, restricted to grey-matter voxels defined by the corresponding tissue probability map [15]). These regional magnitude vectors were then transformed into region–region relationship matrices by computing their outer products, expressing the extent to which regional amplitudes rise or fall together within the component.

**Figure 8:**
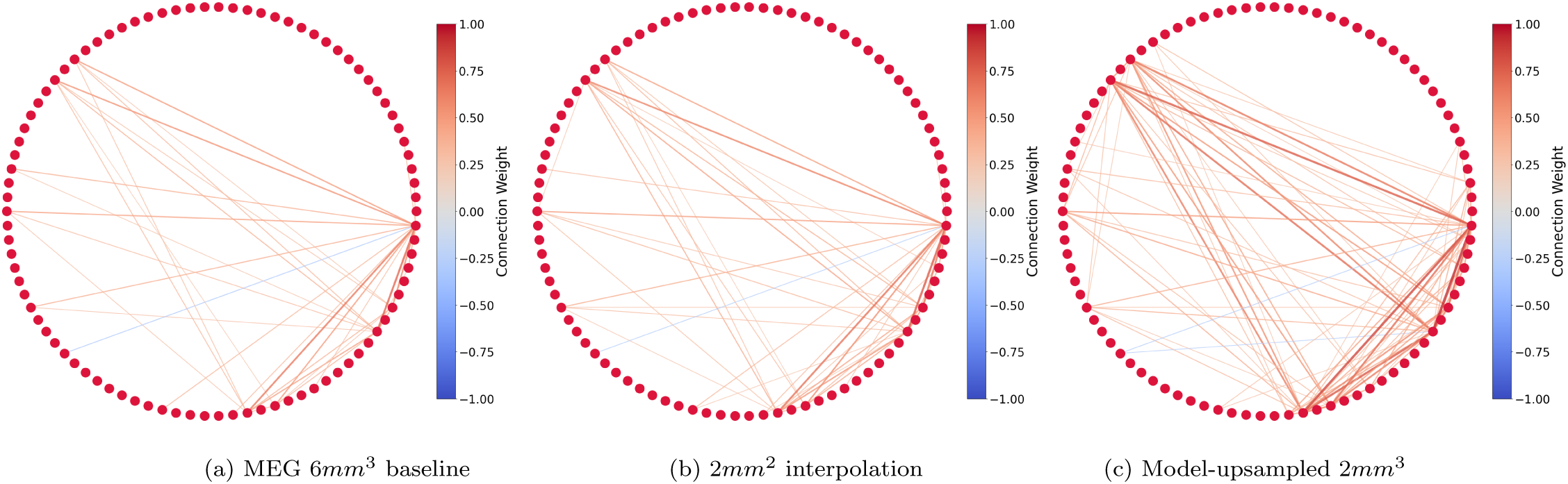
Static region-region relationship, thresholded at |*weight*|=.25; **(A)** Connectome generated at (MEG-native) 6mm² **(B)** Static region-region relationship in maps upsampled to 2mm², as achieved by trilinear interpolation **(C)** Static region-region relationship in maps upsampled to 2mm², as achieved by our BOLD-fMRI-informed model

Region-region similarity patterns show that during the interpolation, we seem to lose salient information (See Fig. 8a and 8b), which is not only re-introduced, but also reinforced in our generated upsampled maps (Fig. 8c). When we calculate average % change of correlations between the regions as a result of the upsampling method, this is supported by overall slight decrease in correlations after interpolation (-0.72%), and overall average increase (15.65%) after modeling.

### 4.3. Emergent detail in upsampled MEG

To assess whether the emergent detail in the upsampled MEG components reflected trans-ferred information form fMRI or distortions emerging from the non-linear warp to standard space, we computed pairwise spatial correlations across all component pairs in a) interpolated, and b) modelled datasets. Similarly, to see if there is a measurable increase in similarity to fMRI, we performed correlations between the modelled/interpolated component, and all fMRI component data. Here, we hypothesize, that if modelling introduced information encoded in fMRI, but otherwise absent in MEG, there should be a measurable increase in overall absolute average correlation coefficients in both.

Both the predicted upsampled and interpolated MEG components exhibited low inter-component similarity (mean |*r*| = 0.054 and 0.023, respectively), indicating preservation of component distinctiveness. Correspondingly, the comparisons to BOLD-fMRI data show a similar, small increase in mean absolute correlation coefficients in modeled (|*r*| = .0879) than in the interpolated components (|*r*| = .0812). While registration to MNI space likely introduces small amounts of information in the data, the ICA extraction which should de-correlate the components as they are being extracted. We believe that the emergence of sulcal and gyral detail in the upsampled MEG maps (Fig. 5) cannot be attributed solely to template bias, as evidenced by the absence of such features in original data, as well as in interpolated controls, but likely reflects the introduction of structured spatial patterns informed by BOLD-fMRI priors. This, along with the modestly higher inter-component correlations in modeled data than in interpolation supports the interpretation that our upsampling approach enhances spatial detail in a meaningful manner by transferring information from BOLD-fMRI data.

### 4.4. Model performance with de-noised data

For this experiment, we individually appraised the extracted 30 BOLD-fMRI components and identified 7 noise components following guidelines outlined in Griffanti et. al (2017) [18]. We then re-trained the model with 1, 3, and 7 excluded noise components selected at random. Results indicate that exclusion of noise does not substantially improve the performance from original *R*^2^ = .92, with *R*^2^ = .92, .92, .91 for 1, 3, and 7 excluded noise components. Results can be seen in Figure B1 in Supplementary materials.

### 4.5. Impact of incomplete data

Through our SHAP analysis, we observed that certain models were predominantly influenced by a relatively small subset of variables (in this case, fMRI components), such as the case with component low-*γ* 18. To investigate the impact of these highly influential predictors and assess how their removal affects the method’s ability to predict and up-sample MEG functional maps, we conducted a retraining process with the top 4 most predictive features for low-*γ* 18 excluded (Figure B2 in Supplementary materials). The results indicate that while the performance is slightly worse, the model is able to substitute missing strong predictors with other, less predictive features which contain similar information.

Next, we excluded components highly correlated to the target component, removing 8 correlates with *r* ≥ .1. Similarly, while doing so negatively impacted performance, the drop was not too pronounced (from *R*^2^ = .92, MSE = .030; SSI = .92 with full data, to *R*^2^ =.88, MSE = .042; SSI = .90 with 8 high correlates dropped) the model was still able to identify other predictive relationships between the BOLD-fMRI and MEG signals. Results can be seen in Figure B3 in Supplementary materials. This finding is encouraging, as it underscores the robustness of the model. Both the performance and the resulting predictions remained relatively stable and high, suggesting that the relationships between the modalities are stronger than indicated by simple correlation analyses.

### 4.6. Half/half validation

Lastly, we decided to test the generalisability of our method between different sets of participants. Considering that models are based on the use of trained voxel-wise relationships between input (30 BOLD-fMRI components) and output (single MEG component), the success of this experiment depends on the similarity of the extracted components across the two halves of our data.

To perform this analysis, we divided the 16 participants into two groups of eight and independently extracted components for each group, obtaining 30 BOLD-fMRI components and 25 MEG components per frequency band, consistent with our primary analysis. The model was trained on data from one group (Group 1) using the previously described procedure and then applied to data from the second group (Group 2).

Given the variability in component order inherent to ICA, it was necessary to harmonize component labels between the two groups to enable modelling. To achieve this, we computed correlation coefficients for each pair of components within their respective groups (e.g., BOLD-fMRI Group 1 vs. BOLD-fMRI Group 2; Alpha Group 1 vs. Alpha Group 2, etc). Components from Group 2 were renamed to correspond to their highest-correlating component in Group 1. If no sufficiently high correlation (a correlation threshold of *r* ≤ 0.4, chosen as a conservative limit to ensure meaningful correspondence) was found between a component in Group 2 and any counterpart in Group 1 , the component in Group 2 was assigned a value of NaN to prevent it from influencing the predictions. Fig. 9 shows results for extracted component most similar (r=.65 for Group 1, and r=.84 for Group 2) to low-*γ*18.

**Figure 9:**
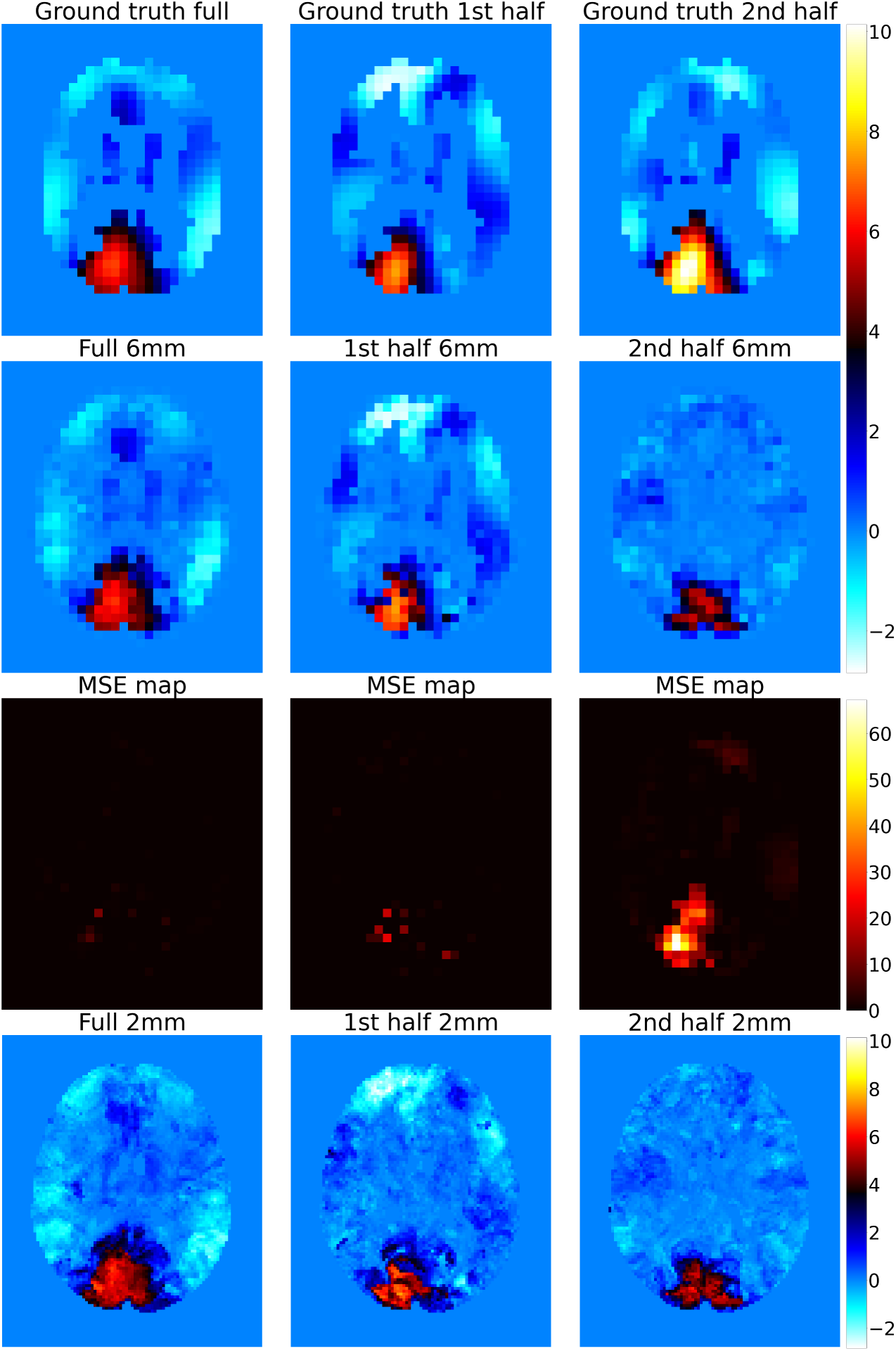
Model performance comparison between model trained on data form **A)** full group-data, **B)** model trained on half of participants, and **C)** pre-trained model from b, with inputs from second half group.

**Figure 10:**
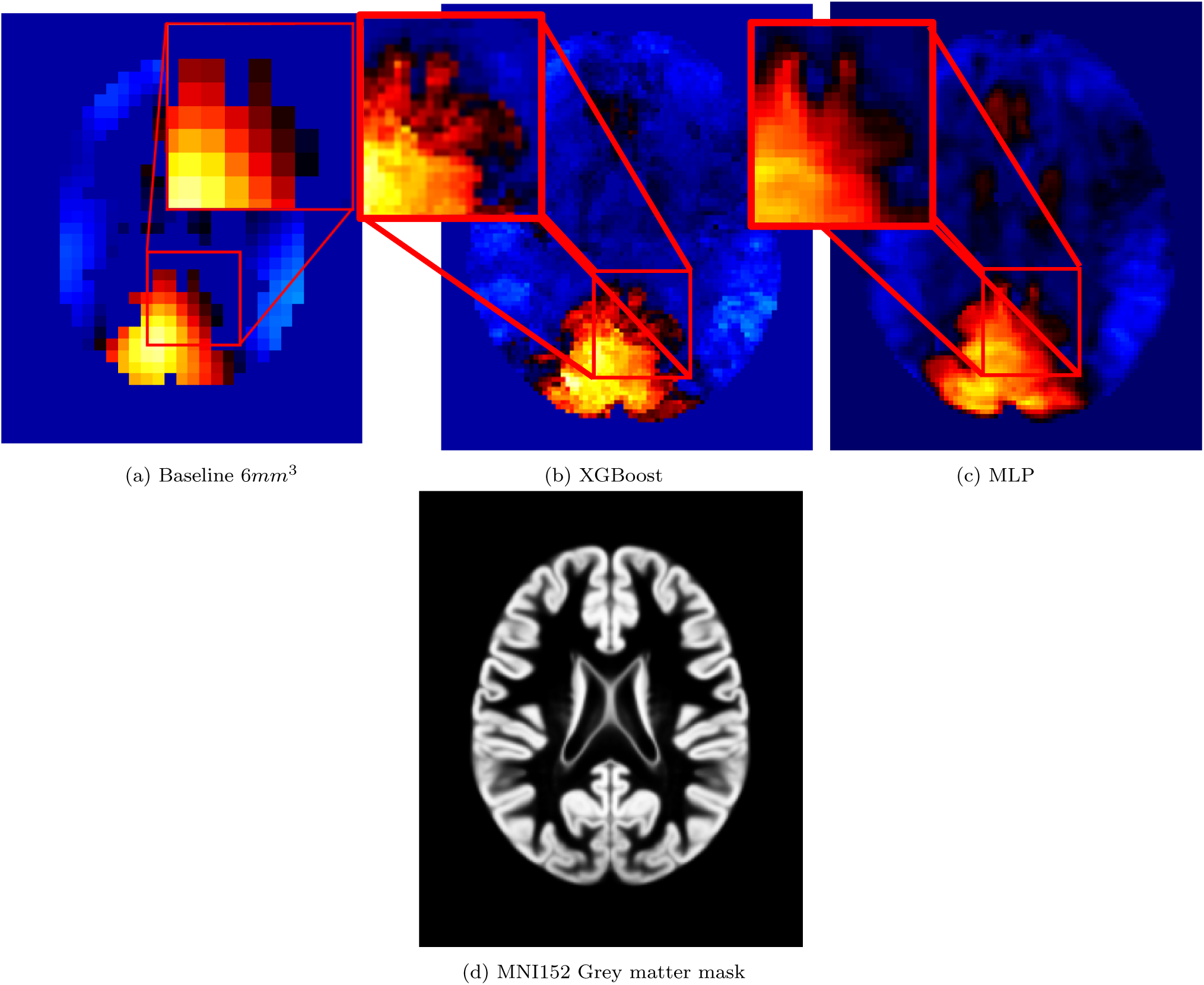
1) Comparison of Early visual network (Component Low-Gamma 18) for (a) 6*mm*^3^ baseline, and super-resolved 2*mm*^3^ maps, as upsampled using (b) XGBoost, and (c) MLP 2) MNI152 Grey matter mask

Visual inspection showed that using data from half the participants successfully reconstructs high-level features, but when compared to maps predicted using a model utilising full set, the predictions become much less consistent, with predicted maps suffering from similar issues as was the case with upsampling *δ*23 in Section 3.2, where predictions were visibly noisy and dissimilar to original maps. As for model’s generalisability between groups, we observed a moderate success, with performance dropping from *R*^2^ = .90 (MSE = .12, SSI = .90) to *R*^2^ = .51 (MSE = 1.16, SSI = .25) between the two halves. This drop in performance can be attributed to the extracted components not aligning perfectly between groups, likely because our sample was too underpowered to allow for sufficient generalisation of independent components between the subsets, with too many individual differences.

### 4.7. From Trees to Neural nets

For our modelling, we have chosen to work with XGBoost for its relative simplicity, good performance, and general explainability thanks to its tree-based nature, which makes it particularly well-suited for finding and subsequently describing latent relationships in the data. Despite this, we cannot ignore existence of more complex architectures, which could be better suited for guided upsampling [35], and have previously even shown to be able to derive latent physical laws without having been trained directly to learn them [55]. To test how much the performance improves using these methods, we applied our pipeline using a multilayer perceptron (MLP), which is often used as a building block of modern deep learning (DL) architectures.

When we assess the results, the MLP models show better performance than our XGBoost model, with *R*^2^ = .93, MSE = .024 & SSI = .83 at 6*mm*^3^ resolution (compared to *R*^2^ = .80; MSE = .030; SSI = .93 in XGBoost), and *R*^2^ = .85, MSE = .051 & SSI = .73 at 2*mm*^3^ (compared to *R*^2^=0.81; MSE = .063; SSI = .86 in XGBoost), respectively. Qualitatively, the upsampled maps are significantly smoother, with more pronounced anatomical detail, clearly showing emergent gyri/sulci. Full MLP performance can be seen in Figure C1 in Supplementary material

## 5. Discussion

This proof-of-concept study tested whether high-resolution MEG spatial maps can be reconstructed from lower-resolution MEG data by transferring spatial information from matched BOLD-fMRI. Preliminary curve-fitting experiments based on classical optimization techniques indicated that without accurate initialisation, reconstructions could achieve similar performance yet remain spatially incoherent, suggesting that while the problem is solvable, it is ill-posed, reinforcing the need for BOLD-fMRI-derived priors to constrain the optimisation. We propose that using explainable machine learning (xML) can effectively solve this issue by learning relationships between the modalities in an explainable manner. The goal of this work was thus, both to enhance MEG’s spatial resolution by utilising quality transfer from paired BOLD-fMRI data, and to examine learned cross-modal relationships, offering a new insight into the neuroelectric–haemodynamic link.

Models trained on 6*mm*^3^ MEG with downsampled BOLD-fMRI predictors generalised well to 2*mm*^3^ BOLD-fMRI input, achieving *R*^2^ = .92 for test subset at 6*mm*^3^, and (as compared to interpolated maps) *R*^2^= 0.79 for a low-*γ* component strongly coupled to early visual cortex. Predictions were evaluated against trilinear interpolation, which served as a reference in the absence of true ground-truth high-resolution MEG data. The model’s outputs had lower MSE and higher SSI, and visual inspection confirmed the absence of hallucinations alongside apparent enhancement of anatomical detail, including sulcal and gyral patterns, potentially linked to vascular anatomy encoded within the (relatively) higher-resolution BOLD signal [56], which, while expected to be encoded within beamformed MEG maps [3], was not visible to the naked eye until upsampled using our fMRI-informed models (Section 4.3). While registration to a shared space likely introduces further shared anatomical framework, this emergence of sulcal and gyral detail in the upsampled MEG maps cannot be attributed solely to template bias. This is evidenced by the absence of such features in interpolated controls, as well as the distinct spatial patterns observed across different modelled components. The model’s reliance on BOLD-fMRI predictors appears to drive a data-informed enhancement of spatial detail, leveraging functional network organization rather than merely reflecting structural alignment.

In our models, SHAP analyses revealed that some, primarily highly correlated MEG–BOLD fMRI components were driven mainly by a small subset of BOLD-fMRI predictors, whereas weaker correlations relied on a more distributed set of predictors (as was evidenced by up-sampling component Delta 23). In SHAP–UMAP space, low-performing models clustered irrespective of frequency band and often corresponded to major networks and/or identified noise components. Further connectomic analyses suggested that interpolation resulted in some network-level organisations disappearing, whereas the model-based reconstructions recovered or more clearly expressed patterns already weakly present in the MEG data.

Performance was robust to input perturbations: removing noise components, top predictors, or high-correlating BOLD-fMRI maps only modestly reduced accuracy (e.g., *R*^2^ drops from 0.92 to 0.88), indicating redundancy in cross-modal spatial correspondences. However, half/half validation revealed reduced generalisability across participant groups (*R*^2^ from 0.90 to 0.51), likely reflecting subject-to-subject component variability — a known limitation of component-based methods in small samples [14].

Following our explainable simple tree-based approach, we performed the modelling using MLP alternative. The model achieved higher accuracy than XGBoost at both resolutions and produced predictions that, on visual inspection, appeared smoother and more anatomically detailed. Another large benefit of using the MLP here is that unlike XGBoost, it considers all features equally, and does not diminish the importance of correlated features, making it more suitable for detailed explanations of inter-modal relationships. However, the reduced interpretability of neural networks compared to tree-based models means that trade-offs remain, particularly when model explanation is a key objective.

A strength of the proposed framework is its fully data-driven nature, avoiding predefined coupling assumptions while remaining interpretable through SHAP. This transparency is further supported by the choice of feature extraction via group-level ICA, which produces spatially and functionally meaningful coherent components. These provide a compact and interpretable basis set for modelling, enabling the learned cross-modal relationships to be mapped onto well-characterised brain networks. The approach operates at the voxel level, maintains low inter-component correlation, and is robust to noisy or incomplete predictors, and the naturalistic viewing paradigm used for data collection improves ecological validity compared to traditional task-, or resting state-based approaches. The approach relies on accurate cross-modal alignment using whole-head source reconstructed MEG data, which has been handled with state of the art tools, guaranteeing minimal bias, however the spatial sensitivity of MEG is highly dependent on SNR [45] which varies greatly between cortical and sub-cortical areas [44, 3]. Limitations include the variability inherent in ICA-derived components, assumption of canonical HRF, which is variable between different networks [41], and the low sample size which likely limited generalisability. Furthermore, the current approach captures only spatial correspondences and does not consider temporal dynamics, underutilising MEG’s temporal richness. It also needs to be stressed that the inter-modal relationships need to be interpreted with caution, as they represent model-level predictivity, where tree-based models, as the one described in this work, can under-emphasize role of highly correlated input features. Finally, while the emergent anatomical features are unlikely to be solely template-driven, residual bias cannot be fully ruled out; however, their absence in the interpolated data strengthens the interpretation that they reflect cross-modal information transfer rather than artifacting.

Overall, our findings indicate that BOLD-fMRI to MEG quality transfer via xML can provide reliable and explainable higher-resolution MEG reconstructions, even when trained on coarser data. The variable performance across components highlights that neurovascular coupling is neither uniform across networks nor frequency bands: strong coupling enables reconstructions from a few key predictors, whereas weaker coupling requires broader integration. The interpretability of the framework offers a data-driven route to probing electro–haemodynamic relationships, accommodating nonlinear dependencies without imposing fixed coupling models.

Future work should integrate temporal dynamics, either using sliding-window or lagged features, or sequential architectures such as LSTMs, recurrent networks, or transformer-based models, to capture evolving multimodal dependencies. This could enhance both spatial and temporal fidelity, enabling dynamic neurovascular coupling analyses under ecologically valid conditions. Improving component stability and expanding datasets will be essential for greater generalisability, and given the variability in SNR in MEG, the best results can be expected in analyses focusing on cortical areas, which would allow for higher effective spatial resolution in the initial source reconstruction of the data.

### 5.1. Conclusion

Using xML, we proposed a novel approach to generate high-resolution MEG maps from BOLD-fMRI inputs and provided insights into the strength and directionality of relationships between the two modalities. This data-guided approach provides an evidence-based method for super-resolution of functional images, and the enhanced explainability of the models provided by SHAP values offers data-driven means of informing the electro-haemodynamic coupling discussion which could overcome limitations of previous literature by allowing for nonlinear description. Furthermore, the proposed method preserves network organisation, introduces biologically plausible spatial detail, and is resilient to noise and partial feature removal.

This work establishes a foundation for a new class of multimodal neuroimaging tools: approaches that use explainable machine learning to transfer high-resolution spatial information from fMRI into MEG without being reliant on model-based biophysical constraints or priors to aid source reconstruction. By demonstrating that cross-modal spatial patterns can be learned, generalised, and interpreted, we show that MEG can gain access to spatial detail previously thought unattainable. This offers a practical, scalable route to improving functional brain mapping, and provides a data-driven lens onto neurovascular coupling itself. Future work incorporating temporal dynamics and larger multimodal datasets has the potential to transform how electrophysiological and haemodynamic signals are jointly understood, ultimately advancing both broader neuroscience and the clinical utility of MEG.

## Authorship Contribution statement

Conceptualization, J.B., M.P., S.K.R., D.M.;

Data Acquisition, P.A, K.D.S.;

Methodology, J.B., M.P., S.K.R., K.D.S;

Validation, J.B.;

Formal analysis, J.B., K.D.S;

Writing—original draft preparation, J.B., M.P., S.K.R.;

Writing—review and editing, all authors;

Supervision, M.P., S.R.;

Funding acquisition, M.P., S.K.R, D.M

All authors have read and agreed to the published version of the manuscript.

## Acknowledgements

We are thankful to Dr. Eirini Messaritaki for consultation in the early phases of the project and Dr. Bethany Routley her invaluable role in data collection.

