## Supplemental materials/Appendix for "Cross-modal quality transfer: enhancing MEG spatial resolution using BOLD-fMRI and Explainable machine learning"

### Supplementary materials: Enhancing brain activity mapping through BOLD-fMRI and MEG data fusion using explainable machine learning

#### A Correlation matrices

##### A.1 BOLD-fMRI x RSN10 Correlation matrix

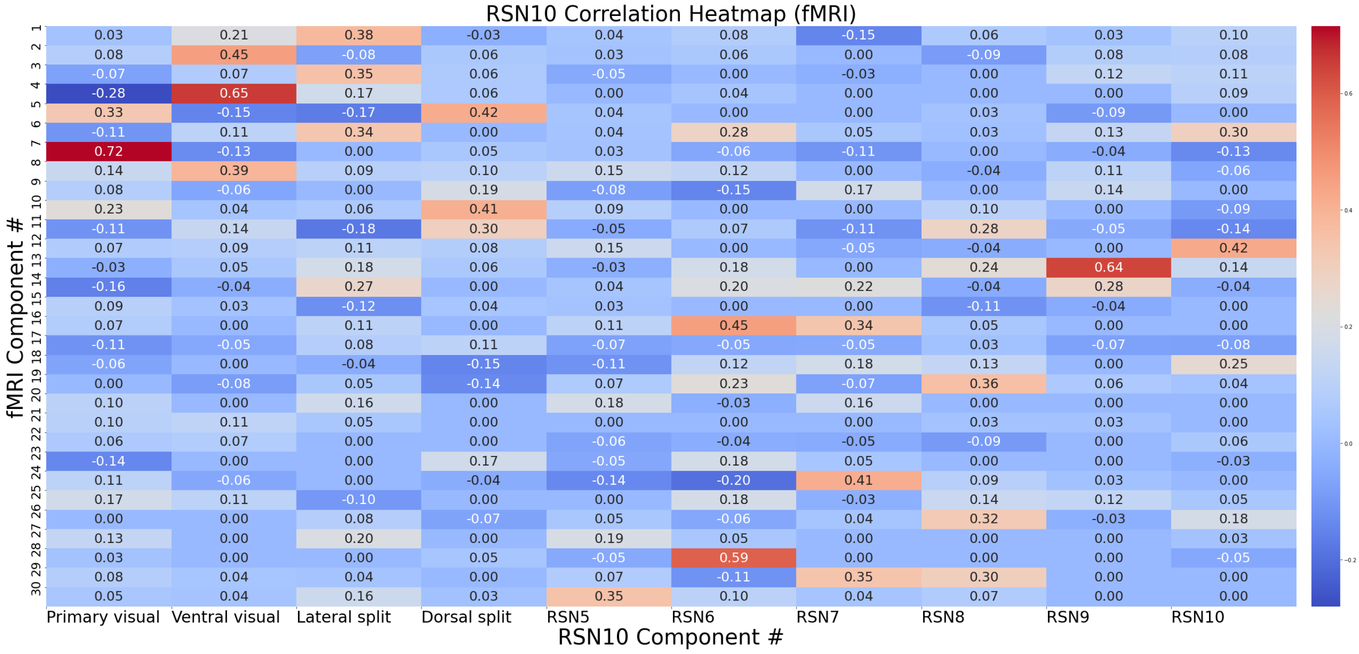


Figure A1: Correspondence between BOLD-fMRI extracted components, and resting state atlas RSN10 as described by Smith et al. (2009) [1]

#### A.2 Correspondence between visual networks and (a) BOLD-fMRI extracted components, and (b) MEG band-specific extracted components

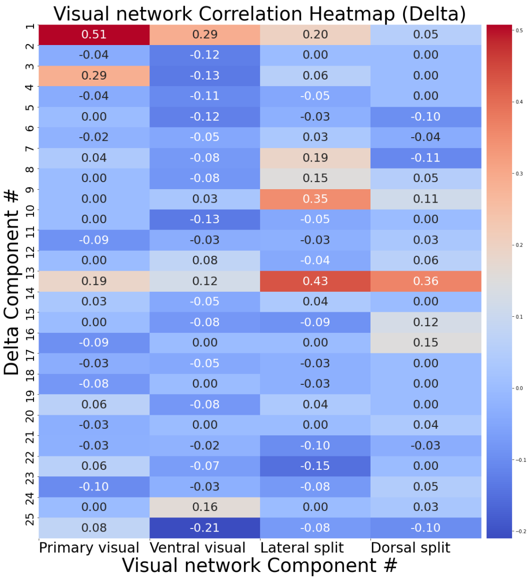


(a) Delta

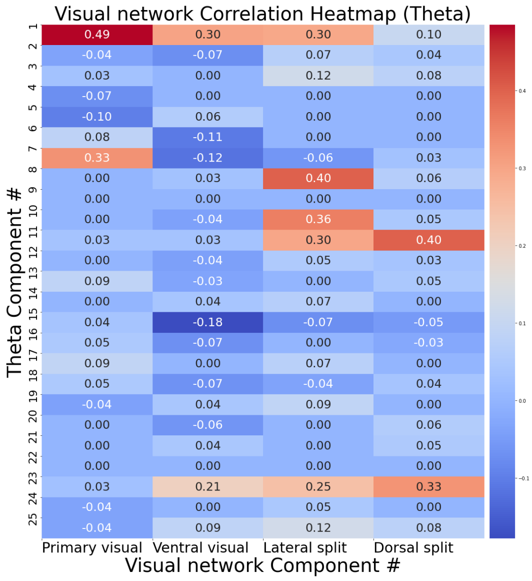


(b) Theta

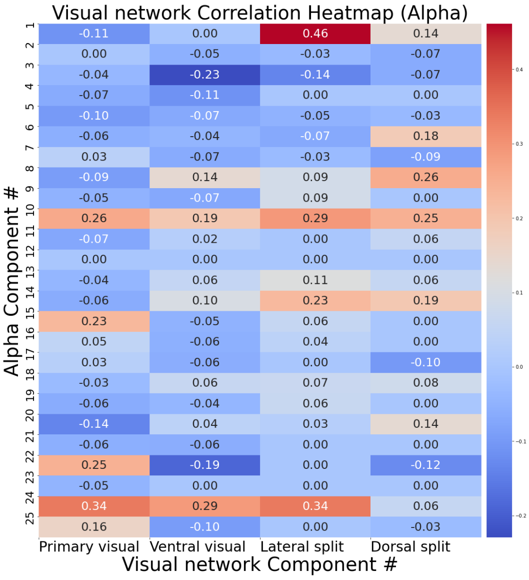


(c) Alpha

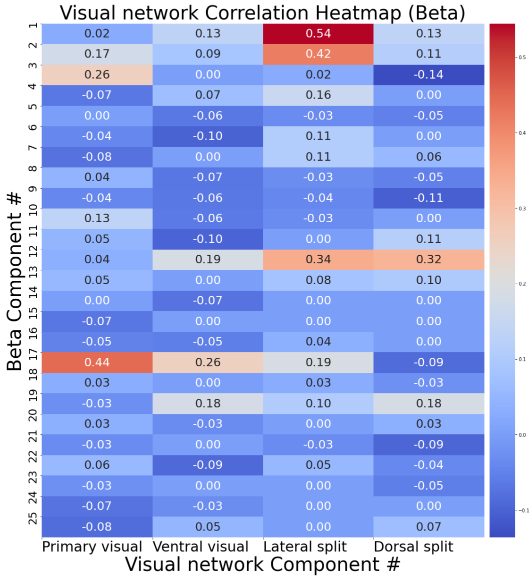


(d) Beta

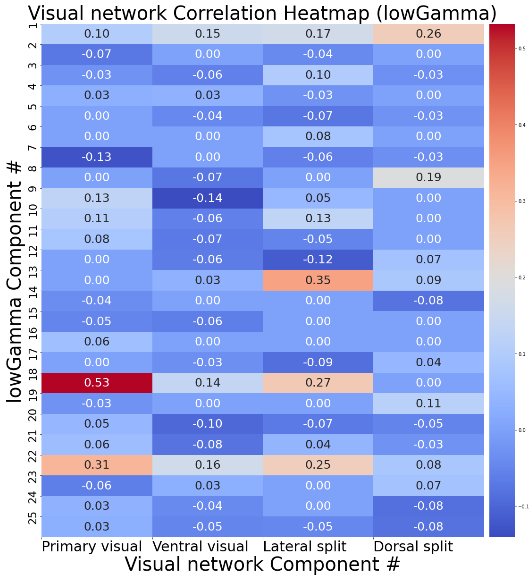


(e) Low Gamma

Figure A2: Visual network correlation matrices for (a) MEG Delta, (b) MEG Theta, (c) MEG Alpha, (d) MEG Beta, and (e) MEG low-Gamma.

### B Incomplete-data models

#### B.1 Effect of denoising

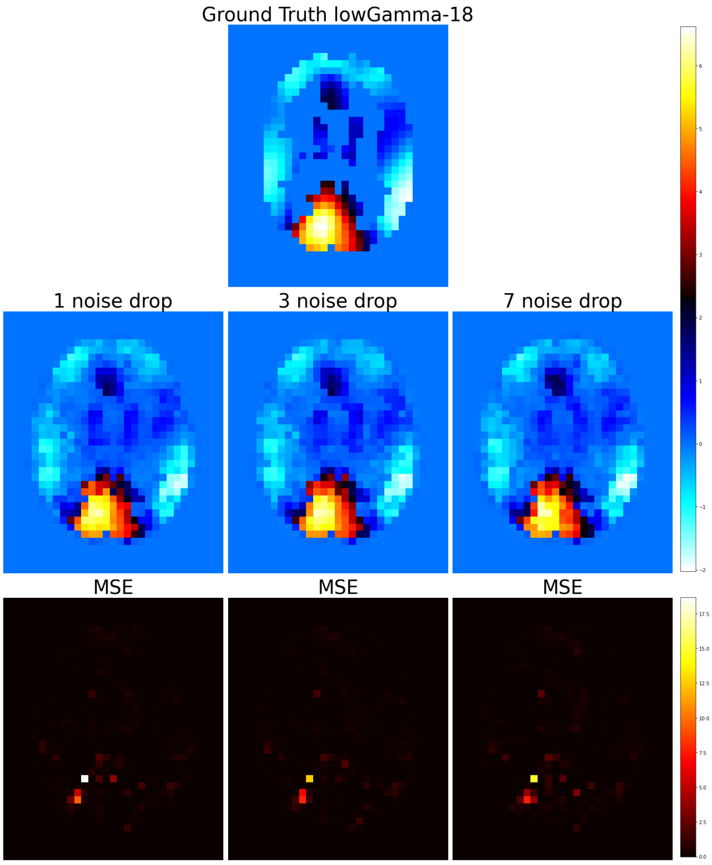


Figure B1: Effect of dropping noise components from the predictor pool.

#### B.2 Effect of dropping top predictors

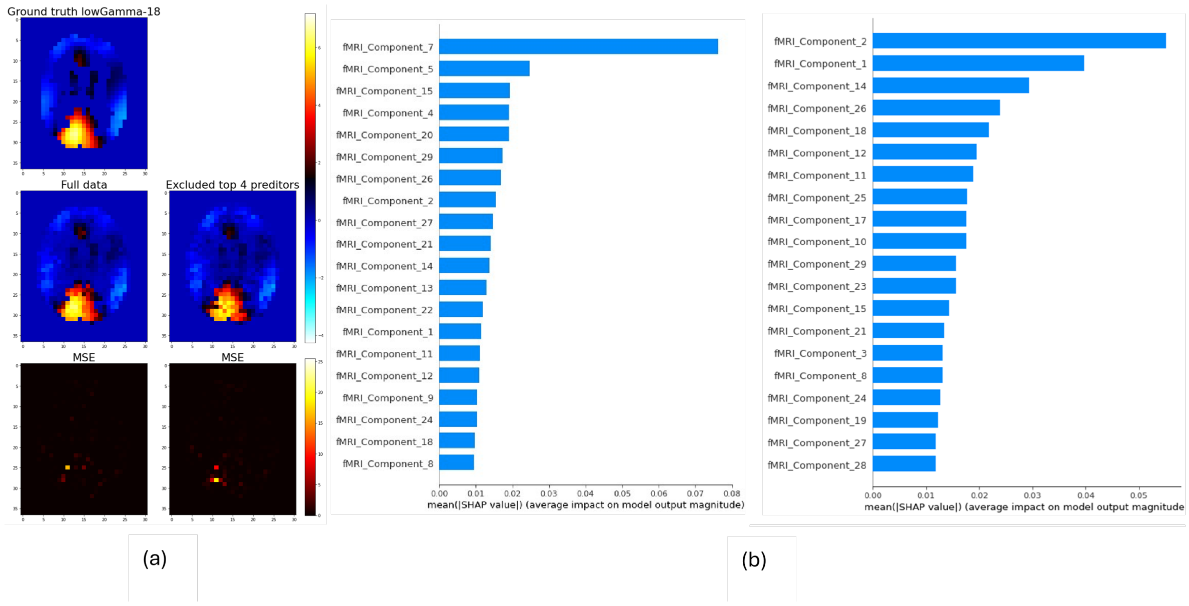


Figure B2: **a)** 1st row: Baseline 6mm map of component low-$\gamma$18, 2nd row: prediction based on intact dataset (right), and on dataset with top 4 predictors missing (right) **b)** shap values for full dataset (left) and for dataset with top 4 components dropped (right)

#### B.3 Effect of designalling (dropping highly correlated with the modelled component)

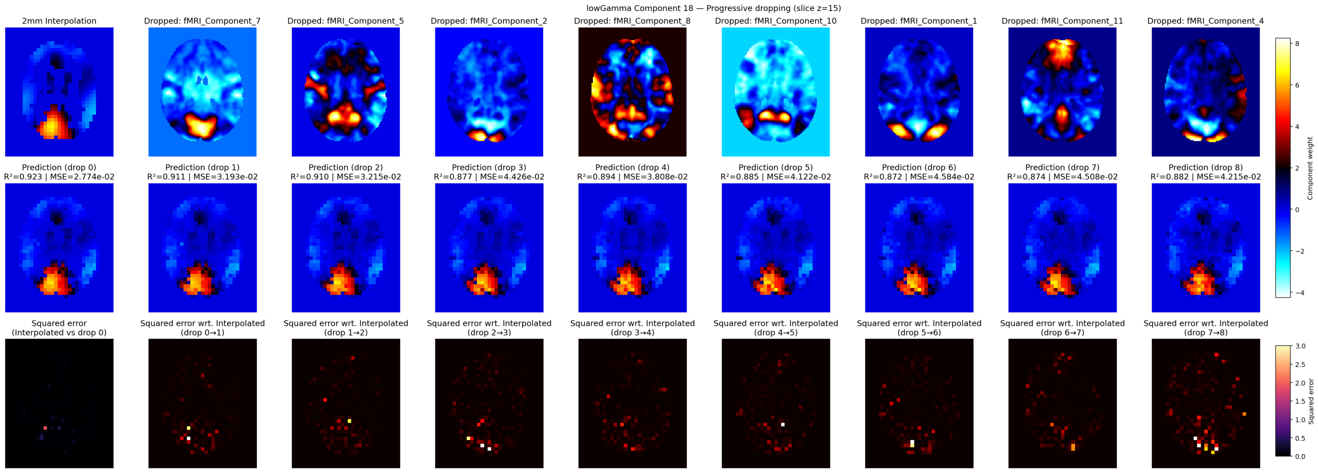


Figure B3: Effect of dropping components highly correlated signal components from the predictor pool. First image on the first row shows modelled MEG component, with subsequent images showing dropped fMRI predictors. MSE maps are have been heavily thresholded for better visibility

### C Multilayer perceptron upsampling performance

#### C.1 Full MLP-generated upsampled maps for early visual network low-Gamma 18

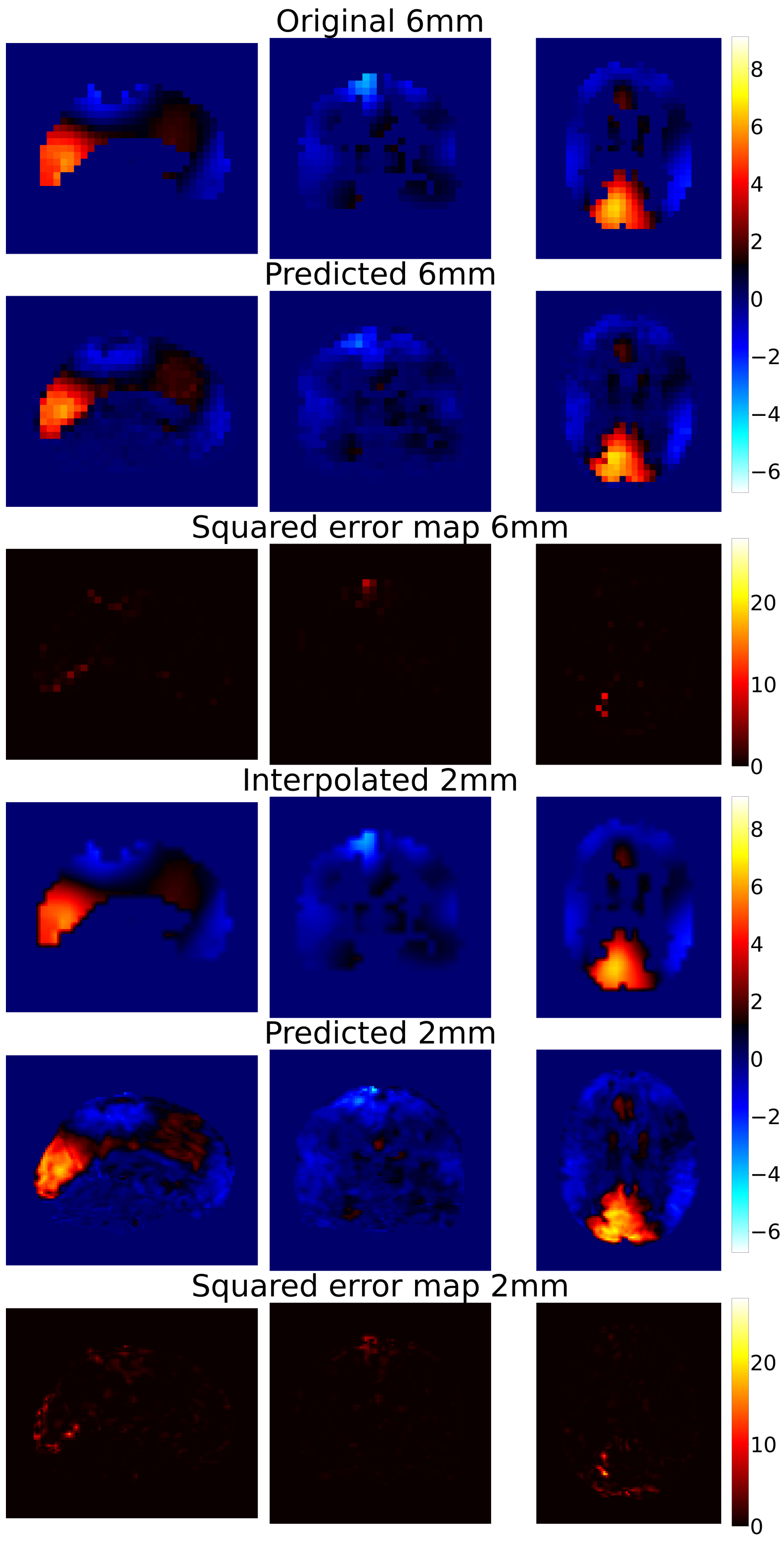


Figure C1: Component 18 from low-$\gamma$ band (40-60Hz). From top: Ground truth $6mm^3$ image, $6mm^3$ image predicted using BOLD-fMRI input using MLP, Squared error maps between original and predicted images, interpolated $2mm^3$, BOLD-fMRI-guided upsampling to $2mm^3$ using MLP
